# Multi-Functionalized Self-Bonding MXene for Minimal-invasive Jet-injected Neural Interface and Tissue Healing

**DOI:** 10.1101/2021.10.18.463745

**Authors:** Baoning Sha, Shengzhuo Zhao, Minling Gu, Guannan Zhao, Liping Wang, Guo-Qiang Bi, Zhanhong Du

## Abstract

Implantable central and peripheral neural interfaces have great potential in treating various nerve injuries and diseases. Still, limitations of surgery trauma, handling inconvenience, and biocompatibility issues of available materials and techniques significantly hinder the peripheral nerve interface for research and clinical purposes. MXenes have great potential as bioelectronics materials for excellent hydrophilicity, conductivity, and biocompatibility. However, their application in bioelectronic interface has been limited due to the poor oxidation stability and fast tissue clearance. Here, we developed a minimal-invasive jet-injected neural interface using MXene nanosheets with strong redox stability, tissue adhesion, conductivity, and good self-bonding properties. We also develop a minimal-invasive jet injector to implant the optimized MXene suspension into the damaged sciatic nerve and establish a neural interface through tissue adhesion and self-bonding. We use this neural interface to promote nerve regeneration and perform electrophysiology recording on moving mice. We prove that the nanosheets can mitigate cellular inflammation, promote tissue healing, and record high-quality electrophysiology signals for predicting joint movement. Thus, our material and implantation strategy together form a novel minimal-invasive neural interface, facilitating the collection and analysis of large-scale living body data to solve the challenge of neurological diseases of the peripheral or even the central nervous system.

## Introduction

Implantable neural interfaces for central and peripheral purposes play an important role in diagnosing and treating various diseases. Now, The Common Fund’s Stimulating Peripheral Activity to Relieve Conditions (SPARC) program is seeking to accelerate the development of maps and tools that identify and influence therapeutic targets—existing within the neural circuitry of a wide range of organs and tissues, especially in peripheral nerves—to improve organ function. This “bioelectronic medicine” strategy could offer new treatment options for diverse diseases and conditions such as hypertension, heart failure, gastrointestinal disorders, and type II diabetes. The mechanisms of these technologies have also been extensively studied and confirmed in recent neuroscience research. However, the peripheral nerve interface’s construction requires surgery, thus increasing tissue damage and risk of inflammation. These factors obstruct the technology to benefit patients. Meanwhile, peripheral nerve injury(PNI) and disorder are among the most common and demanding clinical situations in the neurology field^[1]^. The reason lies in the different causes of nerve injury^[2]^, including (i) cut-off (ii)disease, (iii) stretch-related, (vi) lacerations, and (v) compressions; each demands thorough treatment, especially the latter four. The synergies between the damaged axons, macrophages, Schwann cells, muscles, and surrounding tissues complicate scar formation, preventing effective regeneration^[3]^. Using nanotechnology to regulate cell behavior and escort tissue regeneration becomes the focus of peripheral nerve repair^[4]^. Neural interfaces such as nerve cuffs^[5]^ have effectively repaired severed nerves and restored part of the nerve function. However, placing nerve cuffs requires 360° surgical dissection down to the nerve trunk through skin and muscle, which concomitantly causes more injury. Research has mitigated the tissue response by reducing the stiffness of the interface with conductive self-bonding polymers and nanomaterials^[6]^. Thus, minimizing additional damage and ensuring a healthier tissue-material interaction are the key to improving such techniques.

Since their discovery in 2011, MXenes have flourished as a promising material for high surface area, excellent electrical conductivity, hydrophilicity, mechanical properties, and biocompatibility^[7]^. However, nanomaterials get cleared by cells or diffuse away quickly in vivo, so the current application focuses on the material’s particles properties, facilitating drug release^[8]^, electrode modification ^[9]^ and biosensing^[10]^; furthermore, MXenes oxidize quickly in air and aqueous solution due to large surface area, unsaturated vacancies, and lattice defects^[11]^. These shortcomings obstruct the application of Ti3C2 MXenes, especially in bioelectronics. The demand for functionalized MXenes to tackle tissue diffusion, bio-oxidation, and cell erosion is emerging rapidly. With these challenges handled, MXenes may bloom in implantable neural interfaces and facilitate neuroscience research^[12]^, such as the diagnosis and treatment of neurological diseases by neural recording and stimulation^[13]^, and the enhancement of wearable, flexible sensors to form information networks and record physiological conditions.

Conventional MXenes modifications focus on stability, conductivity, and dispersion separately. Strategies like changing dispersion medium^[14]^ or introducing polyanions like polyphosphates, polyciliates, and L-ascorbates^[15]^ to passivate lattice defects on the edges can increase stability but may reduce conductivity or biocompatibility. In our multifunction-alization approach, we use PDA and PEDOT to passivate the lattice defects of MXenes and increase the in vivo stability. Furthermore, the redox of phenol-quinone in PDA can prevent tissue diffusion with increased adhesion and relieves inflammatory response by scavenging reactive oxygen species (ROS). The introduction of PDA and PEDOT layers also increases the stability, d-spacing of the nanosheets, and the dispersibility in hydrogel. Specially, we adopt an oxidant-free strategy to polymerize the PEDOT with a low polymerization degree. This layer reinforces the self-bonding and film-forming performance of the nanosheets^[16]^. Next, we design a novel minimal-invasive jet-injected neural interface based on the above characteristics, promoting beneficial interactions among Schwann cells, macrophages, and neurons, increasing cell viability and differentiation, and repairing the damaged nerve. Due to the intrinsic oxidative protection capability, this material can benefit drug delivery, especially molecules with redox bioactivity, which often relates to high biological value for disease treatment. Next, we use a deep neural network to analyze the signals obtained with the interface and predict joint motion accurately. We believe this novel jet-injected interface can reliably collect a large amount of data, providing a breakthrough for clinically challenging neurological diseases.

## 2. Results and Discussion

### 2.1 Concept and Multi-functionalization strategy

To form a stable in vivo interface, the oxidation and diffusion of the nanosheets must be reduced. First, we use Na ions to intercalate the Ti3C2 layer and obtain MXene-N; then, we use phenylsulfonic acid diazonium salt for surface modification to obtain MXene-NS. Here, sodium ions increase the carrier density and provide excellent electrochemical performance while weakening the inter-layer cohesive forces, expanding the d-spacing, and accommodating diazonium salts^[17]^. The bonded diazonium salts further expand the inter-layer spacing and delaminate the multilayered MXene-N. PEDOT with various co-dopants such as CNT and ionic liquid had demonstrated outstanding performance for recording and stimulating neural interface electrodes due to the high conductivity and stability of the polymer^[18]^. Thus, we use PEDOT to passivate the lattice defects of Ti3C2 and improve its antioxidant capacity, conductivity, and biocompatibility. However, traditional synthetic processes—like chemicalpolymerization^[17]^, electropolymerization^[19]^, and photopolymeri-zation^[20]^ —are not ideal for coating PEDOT to MXene-NS: the in-situ chemical polymerization usually requires oxidizing catalysts (Fe3+), which will destroy the MXene-NS structure. Meanwhile, electropolymerization and photopolymerization require harsh reaction conditions and can hardly form dispersed nanosheets.

For this reason, we develop an oxidant-free strategy to coat PEDOT on MXene nanosheets. Studies have shown that charge transfer can induce EDOT polymerization between Ti_3_C_2_ layers^[16]^. According to Bader analysis^[21]^, when aggregation occurs, there are 0.34-unit electrons transferred from EDOT to Ti_3_C_2_ due to a higher Fermi energy level aligned to the vacuum energy level of EDOT at −4.31 eV compared to Ti_3_C_2_ at −6.24 eV. However, previous research considered polymerizing EDOT on the accordion-like MXene, rather than the monolayer MXene. For the in-situ polymerization of the stripped monolayer Ti_3_C_2_, the potential electron transfer acceptors density of EDOT are reduced by half, resulting in a decrease in the reaction speed and agglomerating PEDOT to sediment (Figure 3a). Studies have proved that sulfonic acid groups and catechol groups can promote and accelerate EDOT polymerization^[22]^: we polymerize dopamine (DA) on MXene-NS through deprotonation and intermolecular Michael addition reaction to form a cross-linked PDA film, introducing catechol groups. After the above functionalization, EDOT can be polymerized in situ by simply mixing the aqueous solution without using any oxidant, validating that the energy cost of polymerization is reduced along with the charge transfer during the polymerization process. We later confirm that the PEDOT polymerized in this way has a low oxidation level (Figure 3i) and tends to polymerize further, which optimizes the self-bonding and film-forming performance. As shown in Figure 2, the PDA and PEDOT layers can passivate the lattice defects of Ti_3_C_2_ and support better in vivo stability; the electron flow among PEDOT, PDA, and Ti_3_C_2_ activates the redox reaction and increases the oxidation resistance; the redox conversion of phenol groups and quinone groups supports excellent tissue adhesion and reduces tissue diffusion.

Subsequently, we completed the formation of MXene-NSD-PEDOT hydrogel and MXene-NSD-PEDOT jet-injectable dispersion. First, MXene-NSD-PEDOT hydrogel can conduct, record, and output electrical signals. Then, the jet-injectable dispersion of MXene-NSD-PEDOT can penetrate the skin surface to the peripheral nerves and bond into a conductive path by a jet-injector (Figure 1). The material self-bond afterwards and formed neural interfaces for stably transmitting signals between the MXene hydrogel attached to the skin surface to the deep peripheral nerves for the nanosheets’ good oxidation resistance, tissue adhesion, and self-bonding performance.

**Figure 1.**
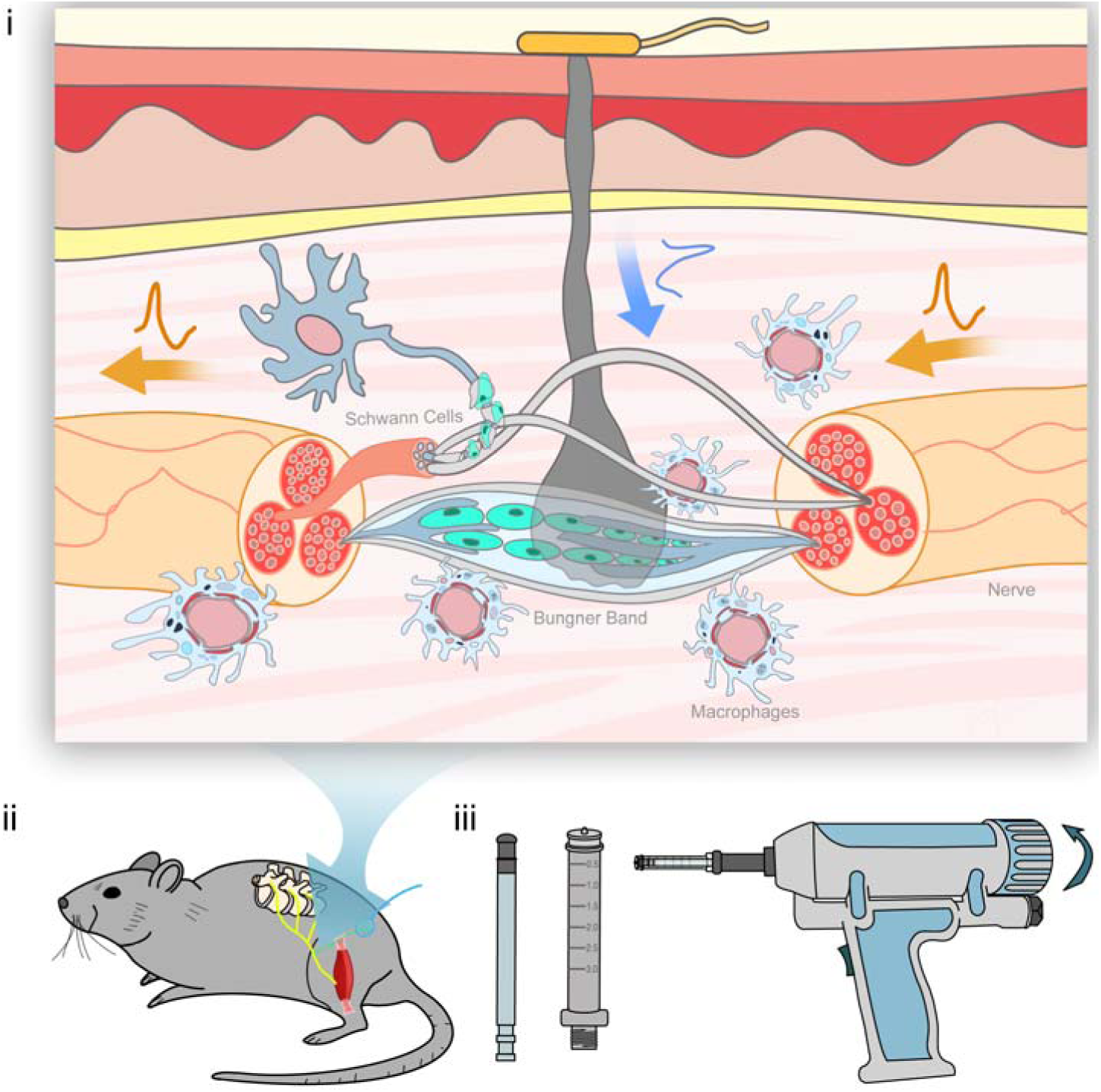
Schematic diagram of jet-injected nerve interface and jet injector. i) Cell interactions with minimal-invasive jet-injected neural interface during tissue healing. ii) Operation design. iii) Customized jet injector.

**Figure 2.**
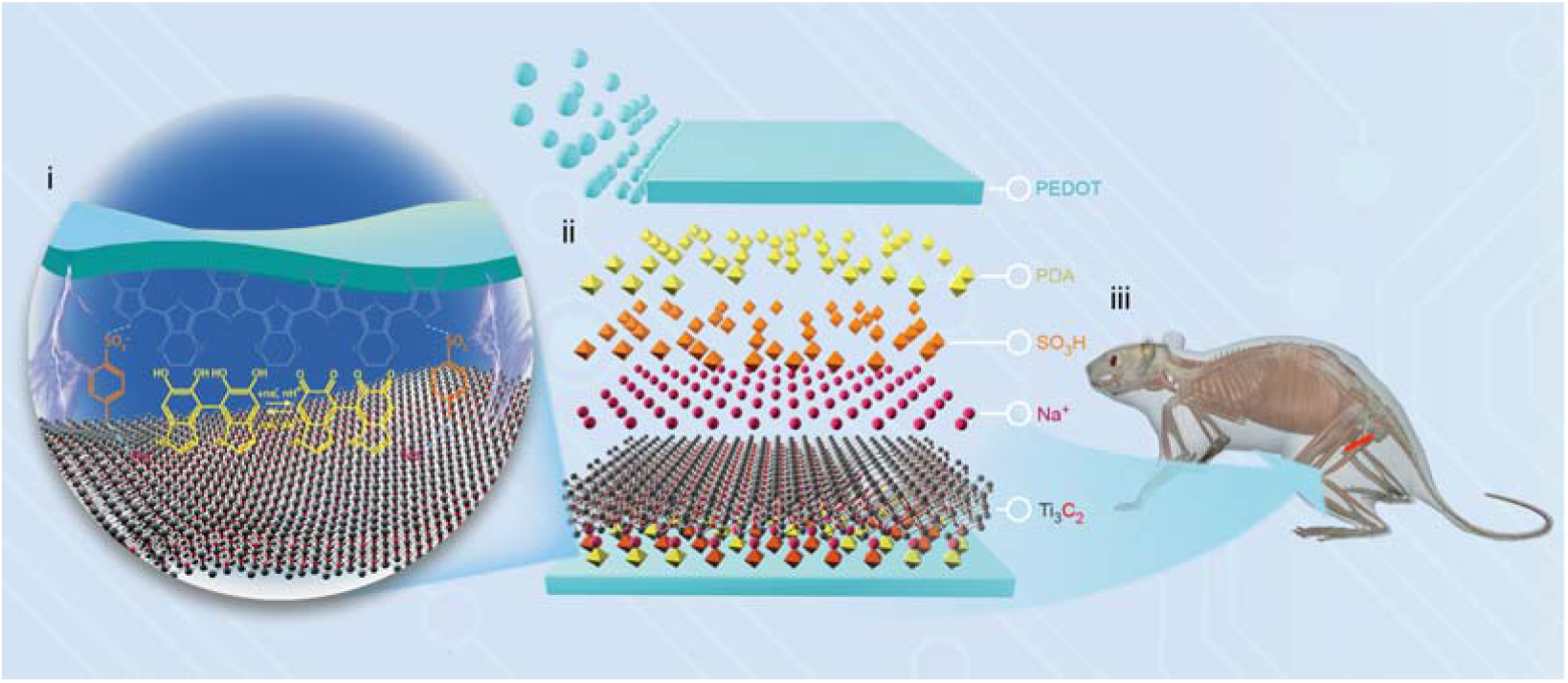
Schematic diagram of MXene-NSD-PEDOT nanosheets. i) The interactions among PEDOT, PDA, phenylsulfonic groups, Na^+^ and Ti_3_C_2_. ii) Formation of MXene-NSD-PEDOT nanosheets. iii) Targeted sciatic nerve.

### 2.2 Characterization of MXene-NSD-PEDOT Nanosheets

First, we use HF acid to selectively etch MAX phase ceramics to obtain accordion-like MXene (Figure 3b) and then use Na^+^ to intercalate MXene and obtain MXene-N. Then we use phenylsulfonic acid diazonium salt to continue the intercalation of MXene-N and obtain MXene-NS. The MXene-NS length is over 100 μm and smooth (Figure 3c). After that, we polymerized a cross-linked PDA film on the surface of MXene-NS through deprotonation and intermolecular Michael addition. The previously modified sulfonic acid groups group can promote the polymerization process of PDA through electron transfer and electronegativity. Therefore, we restrict the thickness of PDA to reduce the aggregation (Figure 3d, i). After the above modification treatment, we mixed MXene-NSD with EDOT ethanol solution to polymerize the PEDOT layer in situ. The synthesized MXene-NSD-PEDOT nanosheets are shown in (Figure 3d), where pink represents the middle layer of MXene-NSD. The thickness of the PDA layer is significantly lower than that of PEDOT (yellow), which increases the stability during the synthesis process and maintains the integrity of the shape.

The synthesized MXene-NSD-PEDOT nanosheets have good self-bonding performance because of the low polymerization degree of PEDOT coating. Figure 3e-f shows the morphology of MXene-NSD-PEDOT dispersion after natural drying in the air: the folds and bumps on the surface enhance the electrical properties and cell adhesion of the nanosheets; the PEDOT layer with low oxidation degree further polymerizes in air condition and form a large area of covalent connection between the nanosheets.

**Figure 3.**
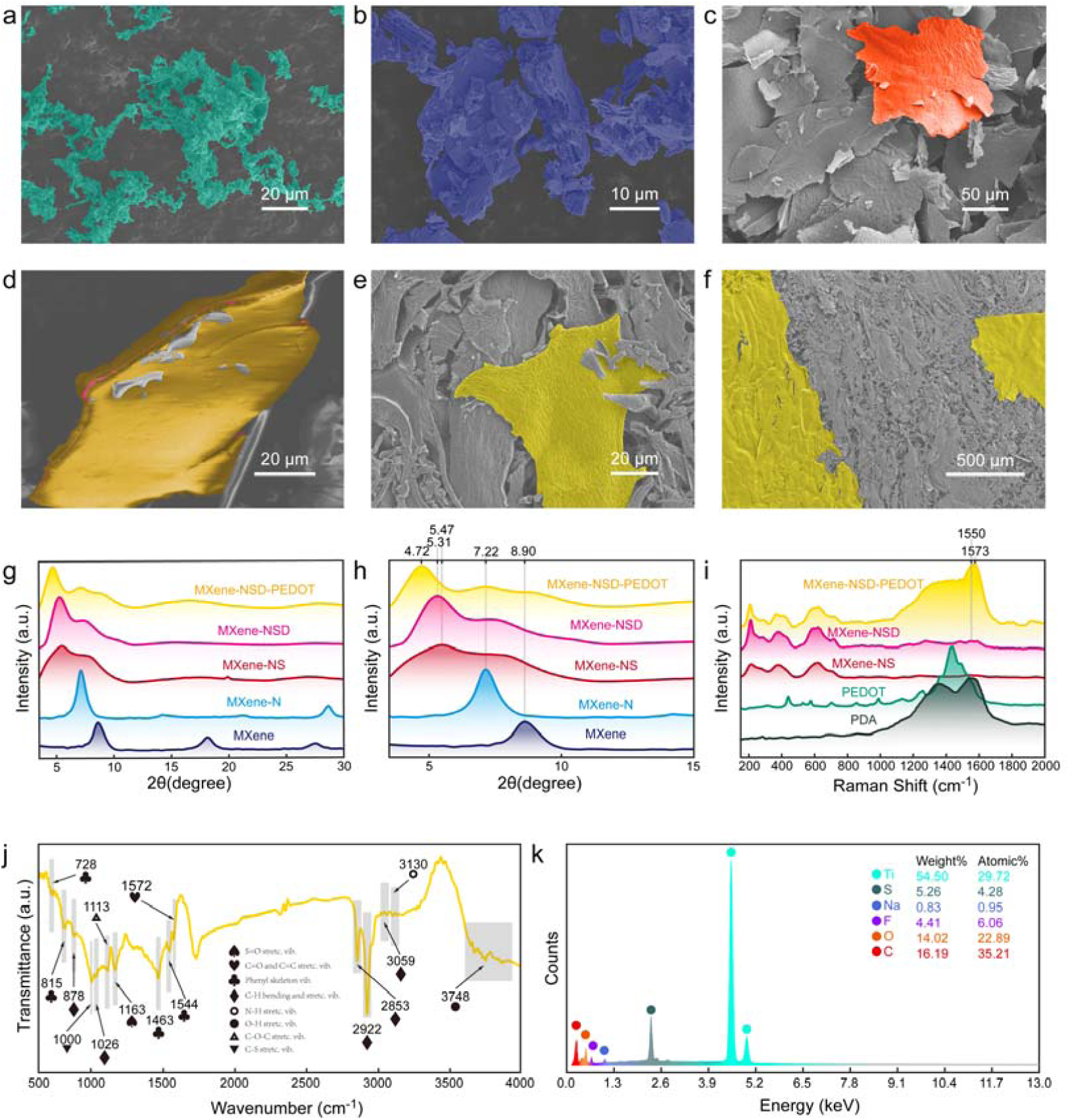
Synthesis of the MXene-NSD-PEDOT nanosheets. SEM micrographs of a) MXene, b) MXene-NS, c) conventional synthesized PEDOT with oxidant, and d-f) MXene-NSD-PEDOT nanosheets. The layer was colored to facilitate visual observation: yellow, PEDOT layer; pink, MXene-NSD layer; red, MXene-NS layer; dark blue, MXene layer. G-h) XRD patterns of MXene, MXene-N, MXene-NS, MXene-NSD, and MXene-NSD-PEDOT. i) Raman spectra of PDA, PEDOT, MXene-NS, MXene-NSD, and MXene-NSD-PEDOT. j) FTIR spectrum of MXene-NSD-PEDOT. k) EDS analysis of MXene-NSD-PEDOT.

The X-ray diffraction (XRD) patterns for MXene nanosheets are used to examine the structural change of different reactions, as shown in Figure 3g, h. The critical feature of XRD patterns is the significant shift of the prominent (002) diffraction peak to low angle direction. After the insertion of Na ions, the (002) peak shifts to the left by 1.68°, corresponding to a d-spacing increase of about 4.6 Å, which indicates that Na ions are inserted between the MXene layers. Subsequent chemical modification of the surface with diazonium salt caused the (002) peak to move 1.75° to the left, and the d-spacing expanded to 32.28 Å. This further weakens the inter-layer cohesive forces between the MXene-N layers, resulting in large scale delamination; the obtained MXene-NS solution has good dispersibility, as shown in Figure 3c. We notice a broad shoulder of (002) peak and a weakening of (004) (006) appear in MXene-NS probably due to disruption of periodicity induced by soluble aromatic compounds, indicating that the stacked fragments for the Ti3C2 sheets are relatively rare. Later, we compare the XRD of MXene-NSD and MXene-NSD-PEDOT to verify the in-situ polymerization of PDA and PEDOT. The (002) peak continues to move to the left and finally reached 4.72°. The corresponding d-spacing was 37.4 Å, and the (004) (006) peaks almost disappear.

Spectroscopic techniques can further confirm the polymerization of the EDOT. Figure 3i shows the Raman spectra of MXene-NS, MXene-S-NSD, MXene-NSD-PEDOT, conventionally synthesized PDA and PEDOT. The spectra of MXene-NSD and MXene-NSD-PEDOT showed the combination of MXene-NS, PDA, and PEDOT, confirming the polymerization of PDA and the oxidant-free polymerization of EDOT. Precisely, the peaks at 1550 cm^−1^ shown in the MXene-NSD spectrum correspond to C=O catechol stretching vibrations of PDA. The band at 1573 cm^−1^ shown in the MXene-NSD-PEDOT spectrum is related to the C=C vibrations, representing the degree of oxidation of the PEDOT structure. When compared with the conventionally polymerized PEDOT, it appeared that EDOT is less oxidized in MXene-NSD-PEDOT; however, sufficient to initiate the doped polymerization of the PEDOT. (This is also confirmed by cyclic voltammetry in Figure 4e.).

**Figure 4.**
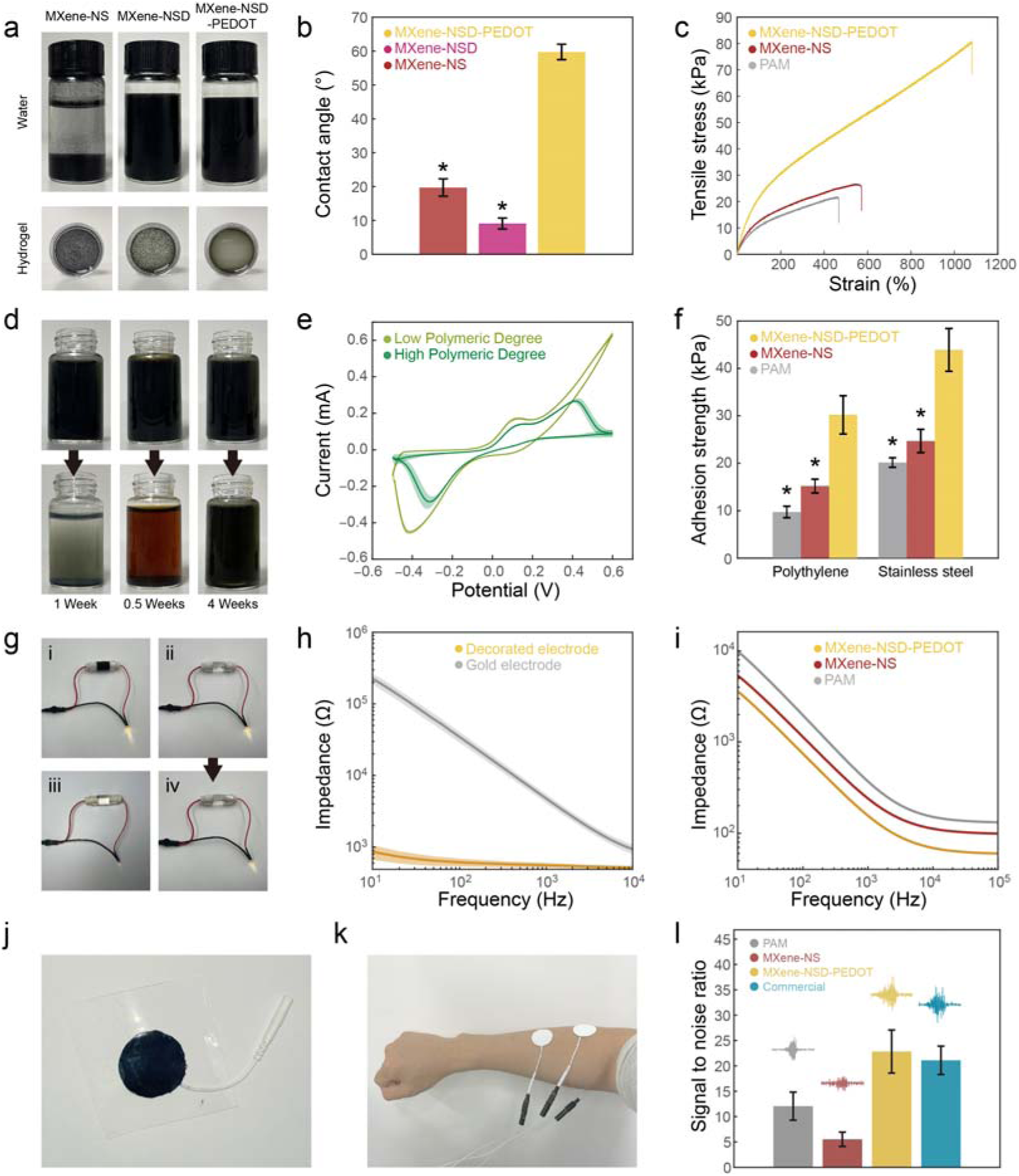
a) Dispersibility of nanosheets in water and PAM hydrogel. b) Contact angle of nanosheets. c) Typical tensile stress–strain curves. d) Oxidation stability performance. e) CV curves of the MXene-NSD-PEDOT nanosheets. f) Adhesive strength of hydrogels to different substrates. g) Powered LED light connected to different materials under a direct current of 3.0 V. h) Impedance curves of gold electrode and MXene-NSD-PEDOT decorated electrode. i) Impedance curves of various hydrogels. * indicates a statistical difference from the MXene-NSD-PEDOT groups. j) MXene-NSD-PEDOT hydrogel electrode. k) Measuring EMG signals with MXene-NSD-PEDOT hydrogel electrode. l) The SNR of EMG signals measured with various electrodes. * indicates a statistical difference from the MXene-NSD-PEDOT groups. (p < 0.05).

The vibration bands in the FTIR spectrum (Figure 3j) of the MXene-NSD-PEDOT at about 3748 cm^−1^ are attributed to the stretching vibration of phenolic O-H. The band at 3130 cm^−1^ comes from the N-H stretching vibration in PDA. The C-H vibration bands are assigned at 3059 cm^−1^, 2922 cm^−1^, 2853 cm^−1^, 1026 cm^−1^, and 878 cm^−1^ (The band at 3059 cm^−1^ comes from C=C-H. The bands at 2922 cm^−1^ and 2853 cm^−1^come from the C-H stretching in PEDOT. The band at 1026 cm^−1^ is attributed to the p-disubstituted phenyl groups in the phenylsulfonic group. The band at 878 cm^−1^ comes from the C-H bending vibration.) The band at 1572 cm^−1^ is assigned to the C=O and C=C stretching vibrations of PDA and PEDOT. The phenyl skeleton vibration is assigned at 1544 cm^−1^ and 1463 cm^−1^. The band at 1163 cm^−1^ is attributed to the S=O stretching vibration in -SO H. The C−O−C stretching bond in PEDOT is assigned at 1113 cm^−1^. The C-S band in the thiophene ring is assigned at 1000 cm^−1^. The bands at 815 cm^−1^ and 728 cm^−1^ are attributed to the P-disubstituted and O-disubstituted phenyl groups in the phenylsulfonic group and PDA. Energy-dispersive spectroscopy (EDS) analysis (Figure 3k) shows the presence of Na, S, and F atoms on MXene-NSD-PEDOT, conforming the previous modification.

### 2.3 Stability, adhesiveness, and conductivity of MXene-NSD-PEDOT nanosheets

The strong inter-layer cohesive forces between MXene nanosheets limits the application in gel modification. Conventional hydrogels usually require special designs like blending graphene^[23]^ or ice mold^[24]^ to ensure system stability. In our design, the phenylsulfonic acid groups increase the electrostatic repulsion between the negatively charged MXene-NS, leading to good dispersibility in water (Figure 4a). We measure the water contact angle of the nanosheets, as shown in Figure 4b. After the polymerization of PDA, the water contact angle decreases from 19.7° to 9.1° degrees, indicating a hydrophilicity improvement. The subsequent polymerization of PEDOT significantly increased the water contact angle to 59.7°. Meanwhile, the dispersibility of MXene-NSD-PEDOT in water remains the same (Figure 4a) due to the combined actions of the hydrophilic groups in MXene, phenylsulfonic acid, and PDA. The MXene-NSD-PEDOT nanosheets hardly undergo aggregation in PAM hydrogel comparing to the MXene-NS-PAM and MXene-NSD-PAM hydrogel (Figure 4a). There are two reasons for this: 1) PDA and PEDOT layers weaken the inter-layer cohesive forces between the nanosheet layers and reduce the aggregation; 2) MXene nanosheets act as cross-linkers and bond in the hydrogel network^[9a]^. These characteristics also increase the mechanical properties of the hydrogels (Figure 4c).

The high electronic activity in MXene-NS accelerates its oxidation. As shown in Figure 4d, the unmodified MXene water dispersion completely oxidizes when exposed to the air for 1.5 weeks, while MXene-NS completely oxidizes within a maximum of 0.5 weeks. Using MXenes in vivo means facing a more severe oxidation challenge because the metabolic activities of cells will continuously produce reactive oxygen species^[25]^. They focus on destroying the crystal defects on the Ti3C2 boundary^[26]^. In our design, the PDA/PEDOT dynamic equilibrium system separates the Ti3C2 crystal from the external environment and induces redox activity by catechol to quinone conversion: 0.34-unit electrons flow from PEDOT to Ti3C2 and reduce the oxidation of catechol group in PDA. We have confirmed an oxidation resistance improvement (Figure 4d) and a six-week in vivo stabilization (Figure 8a iv). Cyclic voltammetry (Figure 4e) is used to investigate the redox behavior of the MXene-NSD-PEDOT nanosheets. In the nanosheets with a low PEDOT polymerization degree, the oxidative peak located at 0.13 V corresponds to the transition of catechol to quinone; the reductive peak located at −0.41 V corresponds to the transition of quinone to catechol. The highly polymerized nanosheets show an additional peak at 0.41 V, which corresponds to the oxidation of the Ti3C2 lattice. Continuing increasing the PEDOT polymerization, we observe a distortion of the CV curve and a sharp rise of impedance. This suggests that as the in vivo time span of the MXene-NSD-PEDOT nanosheets increases, the PEDOT layer will continue to polymerize till a complete state—the electron transfer capability of the nanosheets will reach its peak. From then, the Ti3C2 crystal will participate in the oxidation process: ROS will continuously consume the Ti3C2 electron accumulation and gradually degrade the nanosheet structure. This property may realize a nonlinear degradation of the nanosheets—by accelerating the oxidative decomposition—and a better self-removal ability in the later stage of function.

In nature, mussels can obtain good natural reaction adhesion through continuous redox reactions in a nano-space: Mfp-3 and Mfp-6 form a controlled quinone-catechol group^[27]^. Previous studies have proved that the dynamic redox reaction in PDA can achieve a similar long-term strong adhesion^[28]^. Compared with other two-dimensional materials like graphene, Ti3C2 has higher electronic activity, resulting in higher adhesion (Figure 4f). In contrast, the MXene-NS PAM hydrogel does not show strong adhesion for lacking a redox environment inside the hydrogel network.

As a confirmatory experiment for the subsequent minimal-invasive jet-injected neural interface, we connected different PAM hydrogels to a power supply: the PAM hydrogel with a jet-injected path and the MXene-NSD-PEDOT hydrogel illuminate the LED lights (Figure 4g i-ii). In contrast, the light connected to the uninjected PAM hydrogel is dim (Figure 4g iii). When the jet-injected PAM hydrogel was stretched by 50% fifty times, the LED lamp is slightly dimmer (Figure 4g iv), confirming self-connecting and anti-deformation abilities. Based on these abilities, we hypothesize that the MXene-NSD-PEDOT nanosheets also have an excellent electropolymerization performance. We dispersed the MXene-NSD-PEDOT nanosheets in a PBS solution and deposited the nanosheets on gold electrodes by electropolymerization. The modified electrodes show excellent optical performance and a significant impedance decrease (Figure 4h). We propose that this is because the PEDOT interconnection optimizes the uniformity of the coating. The impedance of MXene-NSD-PEDOT-PAM hydrogel (Figure 4i) is measured: even though the nanosheet content is low, the hydrogel still shows high conductivity. There are three reasons for this: 1) the nanosheets have high conductivity; 2) PDA and PEDOT layers weaken the inter-layer cohesive forces and reduce the aggregation; 3) the nanosheets act as crosslinks in the hydrogel polymerization and form a well-connected network. Therefore, MXene-NSD-PEDOT is a promising material in electrode modification, skin electronics, and neural interface.

As a proof of concept, we validated the advantages of the MXene-NSD-PEDOT hydrogel electrode patches in signal acquisition (Figure 4j). We compared the EMG signals gathered by MXene-NSD-PEDOT hydrogel electrode patches, MXene-NS hydrogel electrode patches, and commercial electrode patches on a human arm (Figure 4k). Compared to the PAM hydrogel electrode, the MXene-NSD-PEDOT hydrogel electrode increases in signal-noise ratio (Figure 4l). In contrast, the MXene-NS hydrogel electrode decreases in signal-noise ratio, suggesting that the MXene-NS nanosheets aggregate in PAM hydrogel and reduce the uniformity and stability of the hydrogel.

### 2.4 MXene-NSD-PEDOT nanosheets scavenge reactive oxygen species

Reactive oxygen species (ROS) are active oxidizing chemicals—including peroxides, superoxides, hydroxyl radicals, singlet oxygen, and α-oxygen—produced during the normal metabolism of cells or tissues^[29]^. The ROS level will rise if the normal physiological metabolism of peripheral tissue is affected by (i) disease, (ii) stretch, (iii) lacerations, and (vi) compressions. Maintaining a low oxidative stress level can coordinate the cleaning function of macrophages and Schwann cells, especially in the first five days after injury. Recent studies have shown that materials with dynamic redox reactions have excellent ROS-scavenging properties^[22]^. In this regard, we investigate the total antioxidant capacity of MXene-NSD-PEDOT with 2,2’-azino-bis(3-ethylbenzthiazoline)-6-sulfonic acid (ABTS): the UV-Vis absorbance of MXene-NSD-PEDOT is less than the absorbance of MXene-NS in the same concentration (Figure 5a), indicating that MXene-NSD-PEDOT has better antioxidant capacity than MXene-NS. Comparing with the Trolox standard sample, we get the Trolox-equivalent antioxidant capacity (TEAC) of MXene-NSD-PEDOT is 10.7 mmol g^−1^, which is higher than the 6.2 mmol g^−1^ of MXene-NS and the 7.4 mmol g^−1^ of MXene-NSD (Figure 5b).

**Figure 5.**
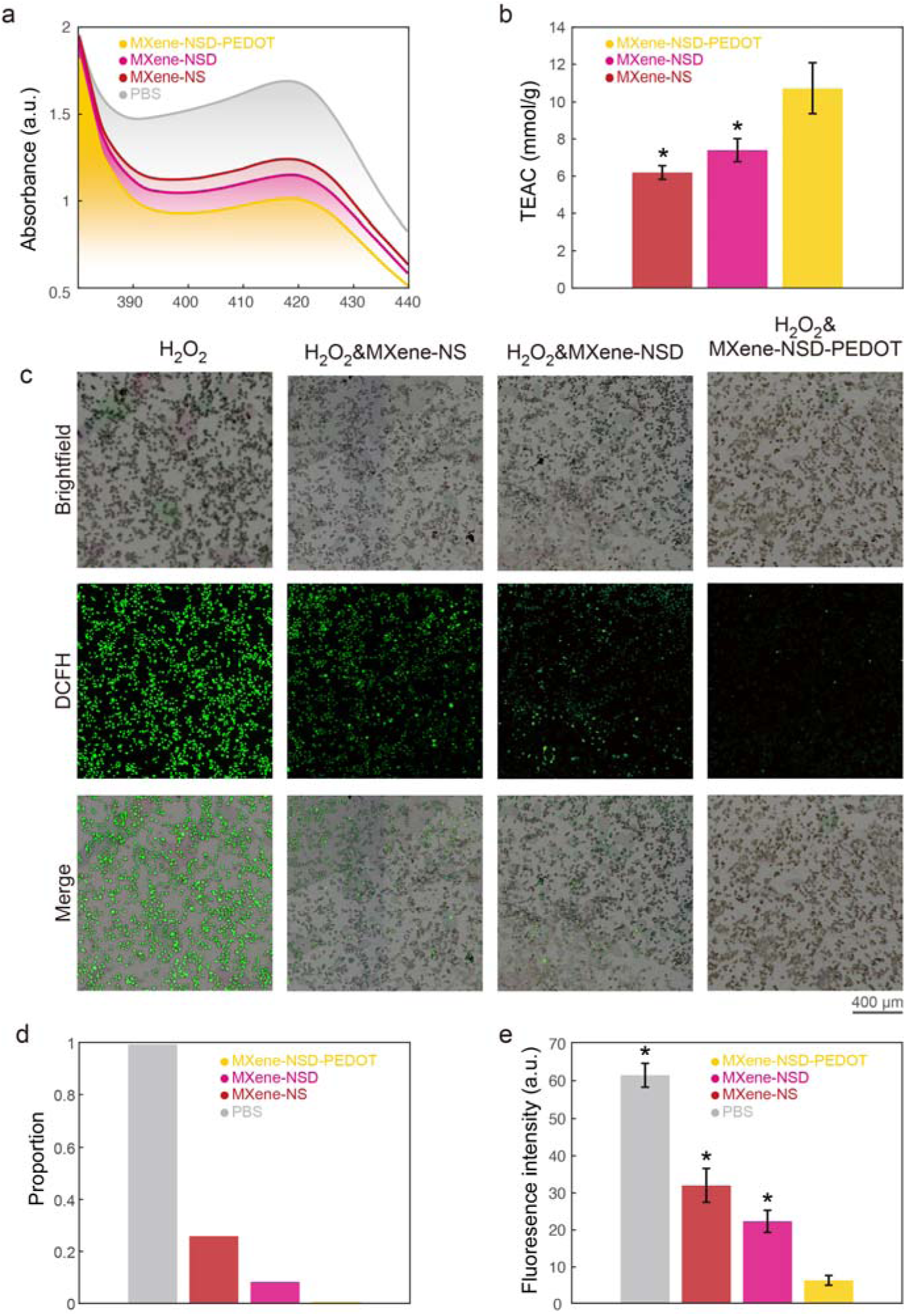
Anti-oxidative properties of MXene-NSD-PEDOT. a) UV−vis spectra of ABTS after reaction with PBS, MXene-NS, MXene-NSD, and MXene-NSD-PEDOT. b) Trolox-equivalent antioxidant capacity (TEAC) of the nanosheets. c) Intracellular ROS-scavenging performance of MXene-NS, MXene-NSD, and MXene-NSD-PEDOT. d) Proportion of cells with significant fluorescence after different MXene treatments. e) Fluorescence intensity of cells after different MXene treatments. * indicates a statistical difference from the MXene-NSD-PEDOT groups. (p < 0.05).

To demonstrate that MXene-NSD-PEDOT can effectively reduce the oxidative stress level of cells during nerve and surrounding tissue damage and coordinate the work of macrophages, we used the DCFH-DA probe to label the Raw264.7 cells stimulated by hydrogen peroxide (Figure 5c). The lowest proportion of cells with significant fluorescence indicates that the cells in the culture medium with MXene-NSD-PEDOT exhibit the lowest oxidative stress (Figure 5d). The fluorescence intensity of the cells in the culture medium with MXene-NSD-PEDOT decrees 5.15 times compared to that with MXene-NS—which is more evident than the 1.7 times increase of the total antioxidant capacity—we speculate that this is because the PEDOT and PDA layer inactivate the nanosheet edges and reduce the cutting stimulation, showing excellent ROS-scavenging properties.

### 2.5 MXene-NSD-PEDOT nanosheets promote Schwann cell migration

In the first five days after injury, the macrophages and Schwann cells (SCs) clean up the axons and myelin debris, leaving empty endoneurium tubes^[30]^. Later, SCs proliferate and transdifferentiate into specialized repair cells, filling the empty endoneurium tubes and forming characteristic Bungner bands or tubes. At the same time, macrophages are also recruited nearby, together with SCs, secreting various neurotrophic factors, which support axonal sprouting and growing toward injured neurons or induce the differentiation of stem cells into neuron-like cells^[4, 31]^. We culture SCs on MXene-NSD-PEDOT and MXene-NS nanosheets to investigate the cell migration properties for nerve repair: we set grid-patterned nanosheets on coverslips in a gap of 0.4 mm; the patterned coverslips are placed into the 24-well plate after UV-alcohol sterilization; we seed the S16 cells(rat Schwann cells) on the coverslips and thoroughly shake the 24-well plate to ensure the uniformity of cell suspension. After two days of culturing, Schwann cells were fixed and stained with phalloidin (green) and DAPI (blue).

As shown in Figure 6a, the SCs growing on MXene-NS is analyzed by comparing the bright field (i) and the fluorescence (ii-iv) images: most of the SCs are round or elliptical; a slight homological distribution of the SCs is observed at the edge of the grid (signs of cell communication and migration). We hypothesize that the excellent conductivity and the abundant attaching sites of MXene-NS induce the communication and the migration of the SCs. Meanwhile, the SCs seeded on MXene-NSD-PEDOT show a clear grid distribution (Figure 6b): more SCs are spindle shape and have extended pseudopods, which connect with adjacent cells and form a network on the nanosheet surface (Figure 6b iii-iv). The ratio of the SCs on the grid pattern to the SCs on the gap of MXene-NSD-PEDOT is larger than that of MXene-NS and increases with the nanosheet concentration (Figure 6c), confirming that MXene-NSD-PEDOT has a better ability to promote cell migration. The average length of SCs on MXene-NSD-PEDOT is longer than that on MXene-NS (Figure 6d), which demonstrated that MXene-NSD-PEDOT could improve the cell morphology and attachment of SCs. Then we test the Viability/Cytotoxicity of SCs on the nanosheets (Figure 6e); the viability is also improved compared with the control group. (This improvement is rare in other two-dimensional nanomaterials like graphene). We propose that PDA and PEDOT can passivate the nanosheet edges, reduce sharp fragments, and prevent cell impairment.

**Figure 6.**
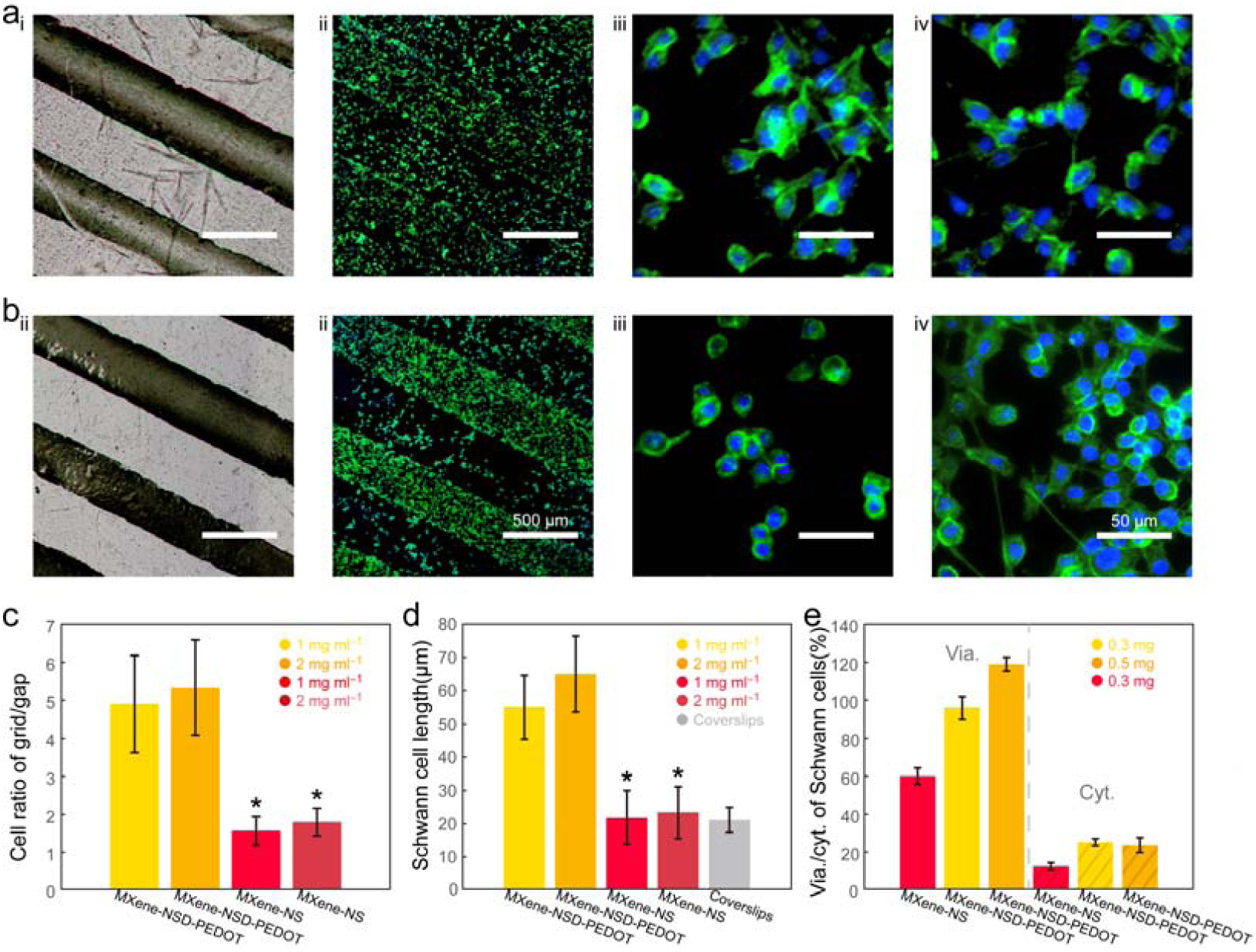
MXene-NSD-PEDOT promotes Schwann cell migration. a) Bright-field (i)and fluorescence (ii) images of SCs cultured on the MXene-NS coverslip (Scale bar: 500 µm); fluorescence images of SCs at the gap (iii) and the grid pattern (iv) (Scale bar: 50 µm) b)Bright-field (i)and fluorescence (ii) images of SCs cultured on the MXene-NSD-PEDOT coverslip (Scale bar: 500 µm); fluorescence images of SCs at the gap (iii) and the grid pattern (iv) (Scale bar: 50 µm). c) Ratio of the cells on the grid to the cells on the gap. d) Average lengths of SCs cultured on MXene-NSD-PEDOT, MXene-NSD-PEDOT, and coverslips. e) Viability and cytotoxicity of SCs measured with Calcein/PI assay. * indicates a statistical difference from the MXene-NSD-PEDOT groups of the same concentration. (p < 0.05).

In summary, MXene-NSD-PEDOT performs well in promoting SCs migration for three reasons: 1) the high charge transfer efficiency of MXene-NSD-PEDOT stimulate the intercellular communication; 2) the PEDOT wrinkles on the surface of MXene-NSD-PEDOT increase the roughness (Figure 3e) and hydrophobicity (Figure 4b); 3) PDA and PEDOT layer reduce the cutting impairment of the nanosheet edges.

### 2.6 Electrical stimulation with MXene-NSD-PEDOT nanosheets promote PC12 cell differentiation

Rat pheochromocytoma (PC12) exhibits characteristics similar to neurons and glial cells in physiological morphology and can develop into neuron-like cells in response to nerve growth factor^[32]^ (NGF) and electrical stimulation (ES)^[33]^. Other two-dimensional materials like graphene have previously been reported to promote cell differentiation and growth by improving cell-to-cell communication and cell-material contact^[31]^. To invest the influence of the jet-injected neural interface in the peripheral nervous system, we select an electrical stimulation (ES) model of PC12 cells.

First, we coat the nanosheets on coverslips and put them into a 24-well plate after sterilizing with UV and alcohol. PC12 cells are seeded and thoroughly shacked to ensure uniformity. After three-day culturing with or without ES (Figure 7a), PC12 cells are fixed and stained with phalloidin (green) and DYPI (blue). The MTT data of PC12 on nanosheets (Figure 7b) shows that the viability of PC12 cells without ES is increased along with the nanosheet mass increase, which is consistent with viability/cytotoxicity data for SCs (Fig. 6f). According to Figure 7c, most cells cultured on coverslips (control group) grew in clusters and appeared small and round without any visible axons; the cells on MXene-NSD-PEDOT show shorter axons, suggesting that the MXene-NSD-PEDOT nanosheets can promote the spreading of PC12 cells. In the ES section, the PC12 cells are cultured for one day and stimulate the cells with different ES (30 mV, 60 mV, 120 mV) for 1.5 h. The spreading of ES-treated cells is improved (Figure 7c) compared with cells not exposed to ES; moreover, the cells increase in cell number, axonal number, and axonal length with increasing voltage. Some PC12 cells cultured on MXene-NSD-PEDOT develop axons nearly 100 µm long with 120 mV ES. The proportion of neurite-bearing cells and their axonal length were measured and statistically analyzed (Figure 7d-e)—both are higher in MXene-NSD-PEDOT groups and increase after ES—showing signs of differentiation. These results indicate that MXene-NSD-PEDOT can increase the viability of PC12 cells and induce their spreading and differentiation, which suggests that MXene-NSD-PEDOT is a promising conductive biocompatible material for in vivo bioelectronic applications.

**Figure 7.**
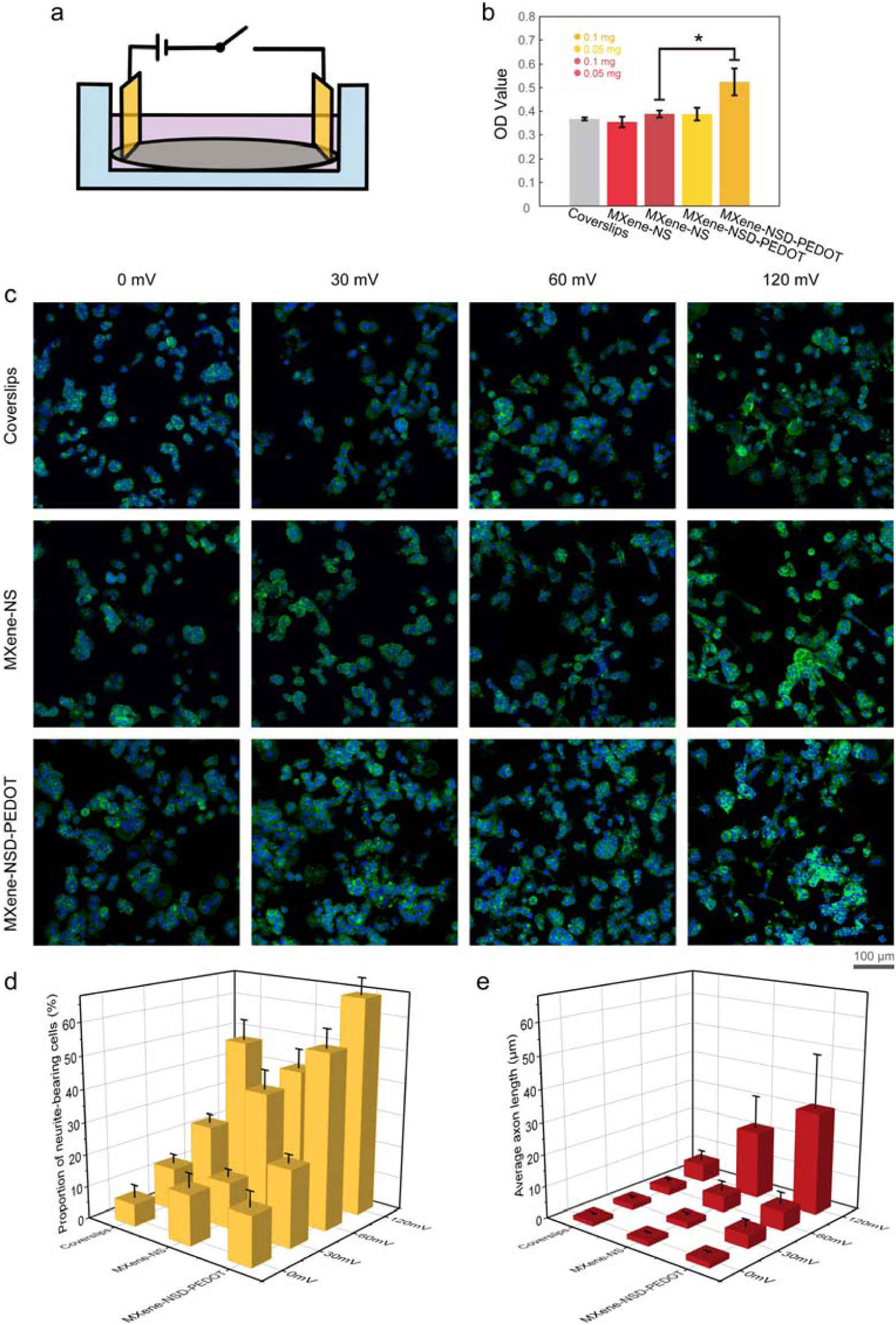
MXene-NSD-PEDOT promotes PC12 cell proliferation and differentiation. a) Schematic representation of the cell culture device with ES. b) MTT results for PC12 cells cultured on coverslips, MXene-Ns, and MXene-NSD-PEDOT. C) Fluorescence images, d) proportion of neurite-bearing cells, and e) average axon length of PC12 cells under different conditions. * indicates a statistical difference from the MXene-NSD-PEDOT groups of the same concentration. (p < 0.05).

### 2.7 Minimal-invasive jet-injected neural interface

The minimal-invasive jet-injected interface consists of two parts; one is the MXene-NSD-PEDOT-PAM hydrogel electrode patches, the other is the MXene-NSD-PEDOT jet-injected conductive pathway. In the follow-up research, we take sciatic nerve crush models with adult C57BL/6 mice to demonstrate the interface function to small nerves, which is trouble in conventional surgical procedures. All mouse experiments were carried out following protocols approved by the Institutional Animal Care and Use Committees (IACUCs) of the Shenzhen Institutes of Advanced Technology (SIAT), Chinese Academy of Sciences (no. SIAT-IRB-170304-NS-DZH-A0297). After anesthetizing the mice with isoflurane, the legs of mice were shaved and disinfected, then the skin where the sciatic nerve was located was cut with tissue scissors, and then the lower muscle tissue was torn with tweezers to expose the sciatic nerve. At the middle part of the sciatic nerve, press the sciatic nerve with a needle holder with a force of 2 N for 10 seconds, release it for 10 seconds, squeeze it again for 10 seconds, and then suture muscle and skin tissue. The tissue was fixed and stained with hematoxylin and eosin (Figure 8a i); sciatic function index (SFI) (Figure 8b-c), motor nerve conduction velocity (MNCV) (Figure 8d), and sensory nerve conduction velocity (SNCV) (Figure 8e) were measured two days later. The histological sections demonstrated that axons are swollen and in poor arrangement; myelin partially disintegrates, and inflammatory cells infiltrate surrounding tissue. The SFI, MNCV, and SNCV show signs of sciatic nerve damage. Then the MXene-NSD-PEDOT nanosheets were dispersed in PBS solution at the concentration of 5 mg mL^−1^, and a jet injection to the sciatic nerve was performed to implant the conductive pathway. After one day of injection, the mice were anesthetized with isoflurane, and the MXene-NSD-PEDOT-PAM hydrogel electrode patches were attached to the skin of the previous injection site; the mice were electrically stimulated (3V 20 Hz 30min) every other day. Figure 8a ii-iii shows the effect of peripheral nerve recovery with the interface implanted and the interface implanted plus electrically stimulated after 28 days, demonstrating that the minimal-invasive jet-injected interface can induce nerve recovery with ES. Figure 8a iv demonstrates the stability of the crosslinked pathway after six weeks of implantation. The nanosheets form a stable connection in the body, and the boundary between the material and tissue remains clear. We measured the SFI, MNCV, and SNCV to characterize the nerve recovery (Figure 8b-e): SFI suggests the recovery of movement performance; the motor and sensory conduction velocity confirm that the minimal-invasive jet-injected neural interface can promote nerve recovery through ES.

**Figure 8.**
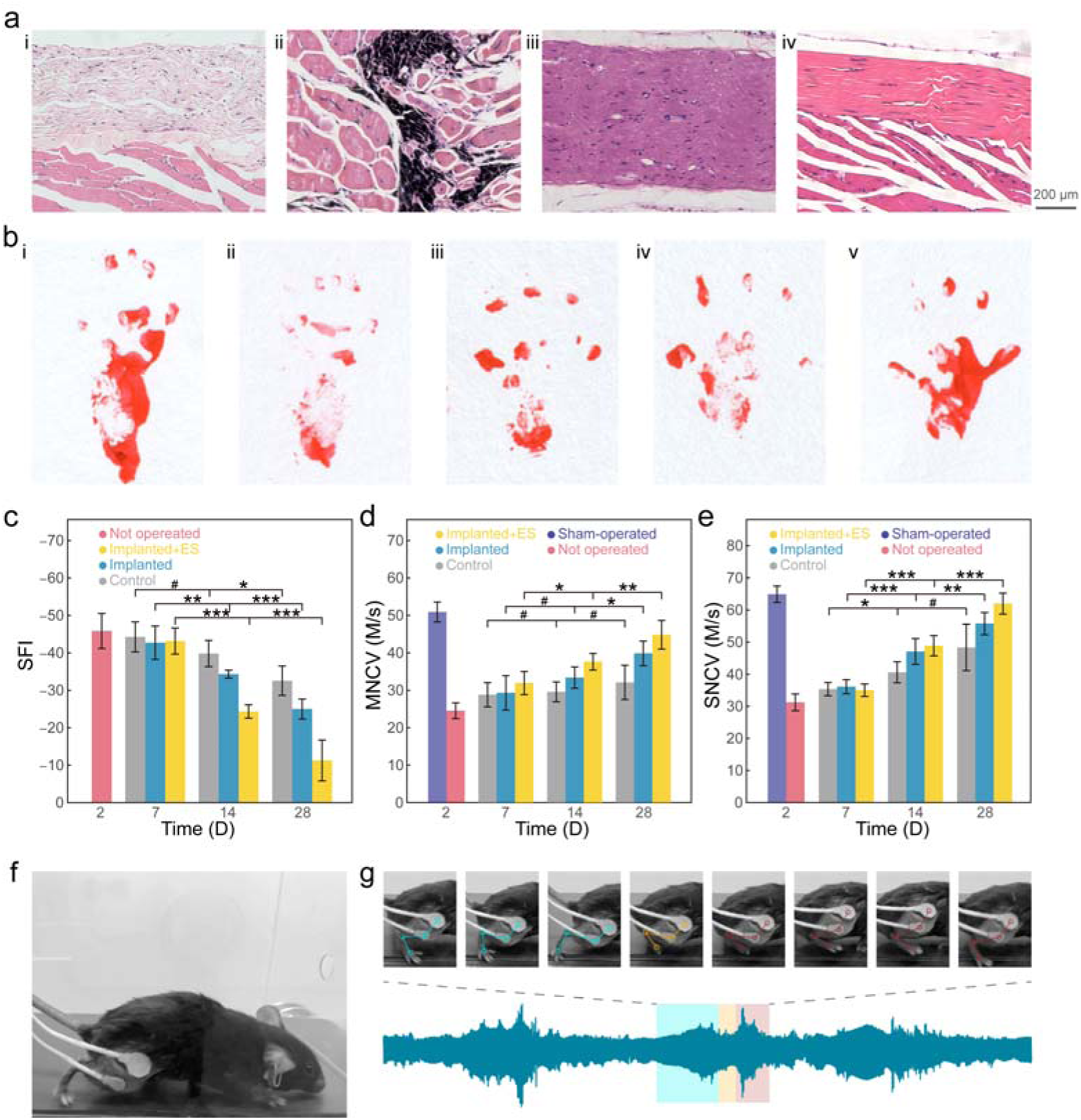
Recovery of sciatic nerve function. a) Cross-sections of sciatic nerves in recovery (i control group, ii interface implanted, iii interface implanted+ES, and iv-six weeks after implantation) and stained with hematoxylin and eosin (HE). b) Footprint stamps in walking track analysis (i 2 days post-operation, ii control, iii interface implanted, iv-interface implanted+ES, and v-sham-operated). c) Sciatic function index (SFI). d) Motor nerve conduction velocity. e) Sensory nerve conduction velocity. f) Photograph of the experimental setup for neural signal recording. g) Synchronous neural signal and joint movement recording. * indicates a statistical difference (p < 0.08); ** indicates a statistical difference (p < 0.05); *** indicates a statistical difference (p < 0.02); # indicates no statistical difference (p > 0.08).

Minimal-invasive jet-injected neural interface is helpful for small nerves that are difficult to record through conventional electrodes. We collected nerve signals from living mice and input the original data into DNN to predict joint movement during exercise. Such prediction is beneficial for diagnosing and rehabilitating nervous system diseases and developing intelligent neural prostheses. In order to obtain the input data of DNN, we recorded the neural signals of mice walking on a treadmill. The treadmill runs at various speeds (Figure 8f). We divide a complete movement into different stages after training DNN with original nerve signals and corresponding joint movements (Figure 8g). The joint movements are predicted by distinguishing neural signals at different stages. Similarly, we also did a simple fitting of the signals collected by the patch electrode. These results show that the minimal-invasive jet-injected interface and DNN can export and process rich signals, predicting joint movement.

## 3. Summary

We realize the stability, biocompatibility, dispersion, viscosity, and self-connectivity modification of Ti3C2 MXene to meet the application needs of the neurological field. Our strategy first passivates the lattice defects on Ti3C2 and strengthens the overall redox activity to strengthen the in vivo stability. The large catechol groups give the material suitable viscosity to reduce the diffusion in peripheral tissue. At the same time, the PEDOT layer with a low polymerization degree gives the MXene-NSD-PEDOT nanosheets good film-forming and self-connection properties. By adding the nanosheets to the PAM network, we developed a tough, adhesive and conductive hydrogel. The nanosheets participate in the polymerization process as crosslinkers. Compared with MXene-NS hydrogel, the introduction of PEDOT increases the d-spacing of nanosheets, resulting in a significant dispersion increase and thus strengthens the mechanical properties and conductivity of the MXene-NSD-PEDOT-PAM hydrogel.

We have developed a new type of peripheral nerve stimulating, recording, and repairing the system based on the conductivity, adhesion, film formation, and biodynamic performance of the MXene-NSD-PEDOT nanosheets. The system uses a jet injector to inject an MXene-NSD-PEDOT dispersion pathway to the peripheral nerve through the skin. The pathway can self-connect and form a stable connection with the help of the adhesive of the PDA layer and the further polymerization of the PEDOT layer. Electrical stimulation can be transferred to the sciatic nerve through PSGO-PEDOT-PAM hydrogel electrodes attached to the skin. It has been proved that MXene-NSD-PEDOT can stimulate the differentiation of nerve cells, maintain ROS stability during trauma, coordinate macrophages’ work, promote Schwann cells’ migration and attachment, and have broad prospects in peripheral nerve recording and wound repair. Then we passed the mouse sciatic nerve trauma experiment to verify the feasibility of this system. Finally, we use the DNN network to model and analyze the large amount of information collected during the wound repair process to fit and predict the activity of the mouse’s legs.

We expect the MXene-NSD-PEDOT nanosheets and the minimal-invasive jet-injected neural interface to be high functioning technologies in neurological disease diagnosis, biosensors, and neural interfaces, which can provide clinically challenging neurological conditions disorders.

## 4. Materials and methods

### Synthesis of MXene-NSD-PEDOT nanosheets

First, 3g Ti3AlC2 powder was slowly immersed into a polytetrafluoroethylene beaker containing 40mL HF aqueous solution and tired at room temperature for 48 hours. The resulting suspension was then transferred to a centrifuge tube and centrifuged at 20,000 rpm for 5 minutes. The wet precipitate was washed with deionized water and centrifuged several times. After decanting the liquid in the last step, 30mL (5 wt%) NaOH solution was added dropwise into the centrifuge tube. The solution was transferred to a beaker and stirred for 2 hours. The product was centrifuged and washed several times with a large amount of deionized water until the pH of the top liquid is 7-8. Then, the obtained MXene-N was redispersed in 10 mL of deionized water for further chemical modification. phenylsulfonic acid diazonium salt were obtained using the literature protocol [38]: 6.3g sulfanilic acid was suspended in 30mL water and was cooled to 0–5°C. A solution of 9mL HCl and 30mL water was pre-cooled to 0-5°C and slowly added to the suspension with stirring under ice bath conditions. After 15 minutes, a cold solution of 2.4g sodium nitrite (18 mL) was added dropwise to the suspension and stirred for 30 minutes to obtain a diazonium salt solution. The diazonium salt solution synthesized above was added dropwise to the MXene-N dispersion in an ice bath with stirring, and the mixture was kept at 0-5°C for about 4 hours. After the reaction, the mixture was centrifuged and washed several times and then centrifuged at 4000 rpm for 1 hour to separate large aggregates and unreacted particles.

Then, the supernatant was lyophilized into the MXene-NS powder. The obtained powder was dispersed with 60 mL deionized water at a concentration of 2.5 mg mL^−1^, and 15 mL Tris-HCl solution (pH 8.5) was added dropwise to the solution. At the same time, 15mg DA was added to 15mL Tris-HCl solution (pH 8.5) and stirred for 15 minutes for pre-polymerization. Then the DA pre-polymerized solution was added dropwise to the MXene-NS solution and stirred for 4 hours to obtain the MXene-NSD nanosheets. The MXene-NSD nanosheets were centrifuged and washed several times and redisperse with 60mL deionized water. Finally, 160 µL EDOT was dissolved in 10 mL ethanol; then the EDOT solution was added dropwise to the MXene-NSD solution and stirred at room temperature for 24 hours. Afterward, the solution was centrifuged and washed several times, and lyophilized to obtain MXene-NSD-PEDOT nanosheets.

### Characterization of the MXene-NSD-PEDOT nanosheets

The morphologies and energy dispersive X-ray spectrum (EDS) of the PSGO-PEDOT nanosheets were examined by scanning electron microscopy (SEM) (MERLIN, ZEISS, Germany) The X-ray diffraction (XRD) was measured with an X-ray diffraction system (SmartLab, Rigaku, Japan). The Raman spectra were measured with a micro-Raman Spectrometer (LabRam HR Visible, Horiba Jobin Yvon, France). The Fourier Transform infrared spectrum was measured with (VERTEX70, Bruker, Germany).

To measure the water contact angle, the freeze-dried MXene NSD PEDOT nanosheets were dispersed in 20 mL deionized water at a concentration of 2 mg mL^−1^. The dispersion was transformed into a self-supporting membrane by vacuum suction filtration. The water contact angle was measured on the self-supporting membrane with a surveying instrument.

To measure Total Antioxidant Capacity of the nanosheets, Peroxidase working solution (20 μL) was added to each detection hole of 96 well plate. Deionized water (10 μL) or the nanosheets dispersion was added into the blank control hole; Trolox standard solution (10 μL) of various concentrations into the standard curve detection hole. ABTS working solution (170 μL) was added into each hole and mixed gently. After 6 min of reacting in dark, the absorption of the samples was measured using a microplate reader (H1M, BioTek, USA)

### Synthesis and characterization of the MXene-NSD-PEDOT-PAM hydrogel

The MXene-NSD-PEDOT nanosheets were dispersed in deionized water at a concentration of 1 mg mL^−1^. 9 mL of the Ti3C2 suspension and 3.0 g acrylamide (AM) was mixed in a glass bottle placed in an ice water bath in Ar atmosphere for 30 min to exclude oxygen gas. 1 mL ammonium persulfate (APS) solution (0.2 mg mL^−1^) 2mg N,N-methylene-bisacrylamide, and 10 μl TMEDA were added into the mixture. Then the mixture was allowed to react 0–5°C under the protection of Ar for 3 min. Finally, the whole mixture was transferred into a mold to form MXene-NSD-PEDOT-PAM hydrogels.

The hydrogels with a bonded area of 25 mm × 25 mm were applied to the surface of the specimens. The samples were pulled to failure at a crosshead speed of 5 mm min^−1^ using a universal testing machine (UTM2203, SUNS, China) until their separation. The mechanical properties of the hydrogels were evaluated using a universal test machine universal testing machine (UTM2203, SUNS, China). Clubbed samples with 6 mm diameter and 60 mm length were prepared. The distance between two clamps was kept at 25 mm and the tensile speed was 10 mm min ^−1^.

### electrochemical characterization of the MXene-NSD-PEDOT nanosheet and hydrogel

The CV measurements of the PSGO-PEDOT nanosheets were carried out on an electrochemical workstation (GAMRY REFERENCE 600, USA). The PSGO-PEDOT nanosheets coated on a gold electrode were used as the working electrode; Pt and Ag/AgCl (KCl sat.) were used as the counter and reference electrodes, respectively. A PBS solution (0.1 M, pH = 7.4) was used as the electrolyte. The group with high polymeric degree PEDOT was electropolymerized under 0.9 V for 70 sec. The potential was varied from −0.5 V to 0.60 V.

The MXene-NSD-PEDOT nanosheets were dispersed in PBS solution (1mg mL^−1^). A gold electrode was used as the working electrode; Pt and Ag/AgCl (KCl sat.) were used as the counter and reference electrodes, respectively. The nanosheets was electropolymerized on the gold electrode under 0.9 V for 40 sec. The impedance of the MXene-NSD-PEDOT nanosheet coated electrode was measured on an electrochemical system (GAMRY REFERENCE 600, USA). The AC voltage was 10 mV rms; The frequency was varied from 10 Hz to 100000 Hz.

The impedance of the hydrogels was measured using a two-electrode setup on an electrochemical system (GAMRY REFERENCE 600, USA). The hydrogels (R = 7 mm, L = 20 mm) were placed between two parallel gold electrodes. The impedance of the MXene-NSD-PEDOT nanosheet coated electrode was measured on an electrochemical system (GAMRY REFERENCE 600, USA). The AC voltage was 10 mV rms; The frequency was varied from 10 Hz to 10000 Hz.

### In vitro ROS-scavenging performance

Raw264.7 cells (6 ×10^4^ cells/sample) were cultured in a 24-well plate with 400 µL of complete medium (90% RPMI 1640 medium, 1% penicillin/streptomycin solution, and 9% FBS). After culturing for 24 h, the groups were washed three times with serum-free medium. DCFH-DA was diluted with serum-free medium at 1:1000 to make the final concentration 10 μ mol L^−1^; and 400 μL of DCFH-DA diluent was added into each group and loaded into cells at 37□ for 30 min. Afterwards, H2O2 (300 mM, 0.4 μL) together with MXene-NS, MXene-NSD or MXene-NSD were added into each group. After 20 min, each group was washed three times with serum-free medium. The groups were observed with a slide scanner (VS120-S-W, OLYMPUS, Japan).

### Schwann cell migration and PC12 cell differentiation characterization

Coverslips were coated with the MXene-NS or MXene-NSD-PEDOT nanosheets in 0.4 mm grid-pattern and were sterilized with 75% ethanol and UV. S16 cells (6 ×10^4^ cells/sample) were cultured on the coverslips in a 24-well plate with 400 µL of complete medium (90% DMEM, 1% penicillin/streptomycin solution, and 9% FBS). After culturing for 48 h, the groups were fixed and stained with phalloidin (green) and DAPI (blue). Schwann cell migration was observed with slide scanner (VS120-S-W, OLYMPUS, Japan). The viability and cytotoxicity of the groups were evaluated using Calcein/PI assay and measured with a microplate reader (H1M, BioTek, USA).

A pair of parallel gold electrodes was used perform the in vitro electrical stimulation (ES). The PC12 cells (6 ×10^4^ cells/sample) were seeded on the MXene-NS and MXene-NSD-PEDOT coated coverslips, respectively, which were sterilized with 75% ethanol and UV. After culturing with complete medium (80% RPMI 1640 medium, 1% penicillin/streptomycin solution, 10% HS, and 9% FBS) for one day, different ES potentials (0, 30, 60, and 120 mV) were applied to each group for 1.5 h, respectively. After 2 days of further culturing, the groups were fixed and stained with phalloidin (green) and DAPI (blue). The groups were observed with a slide scanner (VS120-S-W, OLYMPUS, Japan). The proliferation activities of the cells in the groups were evaluated using the MTT assay and measured with a microplate reader (H1M, BioTek, USA).

### Construction of minimal-invasive jet-injected interface

All mouse experiments were carried out following protocols approved by the Ethics Committee for Animal Research, Shenzhen Institute of Advanced Technology, Chinese Academy of Sciences. The MXene-NSD-PEDOT nanosheets were dispersed in PBS solution at a concentration of 3 mg mL^−1^. Then a jet injector was used to draw 0.3mL of the solution; the knob on the back was adjusted to make sure that the depth of the solution stream reaches the target peripheral nerve. The C57BL/6 mice were anesthetized with isoflurane. The outer thigh of the mice was shaved and disinfected. The mouse’s leg was moved to observe the white sciatic nerve fiber under the skin. Then the nanosheet dispersion was injected to the targeted sciatic nerve with the jet injector. The total injection volume is about 0.05 mL. Later, the MXene-NSD-PEDOT-PAM hydrogel electrodes were attached to the injection site.

### Motor and Sensory Nerve Conduction Velocities Measurement

The Hoffman (H-reflex) was used to determine motor and sensory conduction velocities in large-diameter myelinated axons in the tibial nerve in vivo. The C57BL/6 mice were anesthetized with isoflurane. A pair of needle electrodes were implanted through the surface skin of the proximal metatarsal bone a recording electrode and placed on the foot pad at the root of the thumb of the hind limb as a reference electrode. A ground electrode was placed in the tail. A pair of needle electrodes were implanted to stimulate either the sciatic nerve at the hip or the tibial nerve at the ankle. Square-wave pulses (0.1 to 0.3 msec) were used to stimulate the nerves. Stimulus amplitude was slowly increased until either the H-reflex or M-wave was first discernible. The short-latency M-wave represents muscle electrical activity evoked by direct stimulation of the motor nerves to the plantar muscles. The latencies between the stimulus artifact and the initiation of the M-wave and H-reflex in the muscle were determined for both stimulus locations. The nerve conduction distance was determined by measuring the distance between the most distal stimulating electrode at the ankle and that at the hip after straightening the leg. The motor and sensory conduction velocity were calculated from the following:

Motor conduction velocity = Distance between stimulating electrodes/Latency of M-wave (hip) – Latency of M-wave (ankle)

Sensory conduction velocity = Distance between stimulating electrodes/Latency of H-wave (ankle) – Latency of H-wave (hip)

### Track-Walking movement analysis

the mice were allowed to walk down a wooden walking alley (5.0 * 8.0 * 45 cm^3^). The floor of the alley was covered with white paper. Acrylic paint was applied to the mice’s plantar surface to visualize the footprints. This process was repeated 3 times until clear footmarks were obtained. From the footprints, the following parameters were obtained: distance from the heel to the top of the third toe (print length, PL); distance between the first and the fifth toe (toe spread, TS) and distance from the second to the fourth toe (intermediary toe spread, IT). These measures were made for both the non-operated foot (measurements for this foot are designated as NPL, NTS, and NIT) and the operated, experimental foot (measurements for this foot are designated as EPL, ETS, and EIT). The sciatic function index (SFI) was calculated from the following:

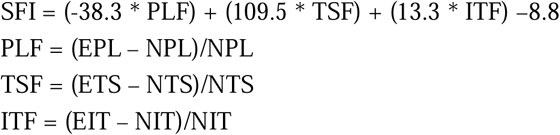

## Acknowledgement

This work was financially supported by Key-Area Research and Development Program of Guangdong Province (2018B030331001, 2018B030338001), Natural Science Foundation of China Grant (31930047, 31700936), National Key R&D Program of China (2017YFC1310503), National Special Support Grant (W02020453), NSFC-Guangdong Joint Fund (U20A6005), the Guangdong Provincial Grant (2017A030310496), Shenzhen Infrastructure for Brain Analysis and Modeling (ZDKJ20190204002), Shenzhen Governmental Basic Research Grants (JCYJ20160531174444711).

